# Proteomic profiling of cytoskeletal interactomes using MT-ID and Act-ID

**DOI:** 10.64898/2026.05.12.724647

**Authors:** Hannah Neiswender, Jessica E. Pride, Rajalakshmi Veeranan-Karmegam, Phylicia Allen, Jamesia Henderson, Mary E. Lowe, Eric A. Vitriol, Kathryn E. Bollinger, Graydon B. Gonsalvez

**Affiliations:** Cellular Biology and Anatomy, Medical College of Georgia, Augusta University, Augusta, GA, 30912, USA; Department of Neuroscience and Regenerative Medicine, Medical College of Georgia, Augusta University, Augusta, GA, 30912, USA

## Abstract

The microtubule and actin cytoskeletons form dynamic, interconnected networks that are critical for eukaryotic cell function. These networks govern intracellular organization, cargo transport, cell migration, and tissue morphogenesis. Microtubules and actin filaments are regulated by diverse binding proteins that control many aspects of their function. However, identifying cytoskeletal-interacting proteins has been challenging due to the transient and weak nature of many interactions and the disruption of native architecture by conventional biochemical approaches. These limitations suggest that numerous physiologically relevant cytoskeletal regulators remain undiscovered. Identifying these factors requires novel and sensitive methodologies that can capture cytoskeletal interactions under native cellular conditions. Here, we present MT-ID and Act-ID, powerful proximity-labeling tools for identifying microtubule and actin-interacting proteins, respectively. MT-ID employs the microtubule-binding domain of MAP7 (EMTB) fused to TurboID, a highly active promiscuous biotin ligase. Act-ID utilizes the actin-binding domain of ITPKA (F-tractin) similarly fused to TurboID. We validate both approaches by successfully identifying numerous known cytoskeletal regulators and discovering potentially novel interacting proteins. Functional characterization reveals that LIMCH1 is a previously unrecognized microtubule-associated protein whose depletion increases microtubule density. Additionally, we identify FBXO30 as a novel actin-interacting protein, with its loss promoting increased focal adhesion formation. MT-ID and Act-ID will be useful not only to identify cytoskeletal interacting proteins but also to define changes to the cytoskeletal interactome when cells are exposed to changing physiological conditions.

## INTRODUCTION

The microtubule and actin cytoskeletons are interconnected and often dynamic networks that are essential to the survival and specialized function of eukaryotic cells. Beyond their roles in maintaining cell shape and rigidity, these networks operate as cellular highways that govern the precise organization of the intracellular environment. Long distance transport of organelles, protein complexes, and mRNA rely on microtubules (1). Conversely, the actin network facilitates short range movement, cortical anchoring, and the generation of propulsive forces that drive cell migration and tissue morphogenesis (1).

The actin and microtubule networks are often quite dynamic and are regulated by a large and diverse set of binding partners. For instance, numerous proteins collectively known as microtubule-associated proteins (MAPs), bind along the lattice of the microtubule and often function to stabilize the filament (2–4). In addition, certain proteins are tasked with nucleating new microtubules whereas other proteins sever existing microtubules into smaller fragments (5–9). Motor proteins of the dynein and kinesin family move cargo along the microtubule track (10–13). Collectively, these factors regulate microtubule growth, shrinkage, organization, and even the activity of motor proteins and cargo transport. In a similar vein, actin filaments are also regulated by a vast array of binding proteins that function in filament nucleation, elongation, branching, crosslinking, and assembly into higher order structures (14, 15). Through the coordinated activity of these binding proteins, actin and microtubules networks can rapidly reorganize during processes such as cell division and in response to signaling cues, developmental pathways, or mechanical forces.

Historically, the identification of cytoskeletal binding proteins has relied on a combination of biochemical purification, genetic screens, and candidate-based approaches. Methods such as co-sedimentation assays, affinity purification, and immunoprecipitation have successfully identified many key regulators of cytoskeletal dynamics (16–21). However, despite their success in identifying cytoskeletal interacting proteins, there are limitations to these approaches. Many cytoskeletal interactions are transient, weak, or dependent on the polymerization state of the filament, making them difficult to capture using conventional approaches (22, 23). Furthermore, purification-based strategies often disrupt the native cytoskeletal architecture by using stabilizing agents or chemical crosslinkers, potentially leading to the loss of physiologically relevant interactions. These challenges suggest that many proteins capable of interacting with actin or microtubules remain unidentified. Novel methodologies that circumvent these limitations are therefore critical for advancing our understanding of cytoskeletal organization and regulation. An additional requirement is the ability to capture these cytoskeletal interactions under physiologically relevant conditions in cells and tissues.

Here, we describe the development and validation of MT-ID and Act-ID as powerful new tools for identifying microtubule and actin interacting proteins. MT-ID involves fusion of the microtubule-binding domain of MAP7 (referred to as EMTB) to TurboID, a highly active promiscuous proximity biotin ligase with a labeling radius of 15 to 30 nm (24). Act-ID uses the actin binding domain of the ITPKA protein (referred to as F-tractin) (25) fused to TurboID. We demonstrate the utility of both approaches in identifying many known microtubule and actin interacting proteins. In addition, numerous potentially novel cytoskeletal interacting proteins were also identified using these approaches. We show that LIMCH1 has a previously unknown role as a microtubule associated protein and that depletion of LIMCH1 results in increased microtubule density. Lastly, we demonstrate that FBXO30 is a novel actin interacting protein. Loss of FBXO30 is associated with increased formation of focal adhesions and internal actin stress fibers.

## RESULTS

### Development and validation of MT-ID

The microtubule-binding domain of MAP7 (referred to as EMTB) fused to GFP, has been widely used to visualize microtubules in cells and tissues without disrupting their dynamics or interfering with vesicle or organelle trafficking (26–28). We therefore reasoned that fusion of EMTB with TurboID, a highly active and promiscuous biotin ligase (24), would enable the in vivo biotin labeling of microtubule proximal proteins (Fig. 1A). Because TurboID has a limited labeling radius of 15-30nm, most of the labeled proteins would likely represent microtubule interacting proteins. We refer to this approach as MT-ID. To test the feasibility of MT-ID, we transfected HeLa cells with a plasmid expressing either TurboID alone or EMTB-TurboID. The cells were then fixed and processed for immunofluorescence. Alexa647 conjugated Streptavidin was used to visualize the localization of biotinylated proteins. TurboID as well as the proteins biotinylated by this enzyme localized diffusely throughout the cell (Fig. 1B). By contrast, EMTB-TurboID efficiently localized on microtubule tracks, as did the proteins biotinylated by this fusion construct (Fig. 1C). Similar results were obtained with other commonly used cell lines such as Cos7, U2OS, and hTERT-RPE (Supplemental Fig. 1A-C).

**Figure 1:**
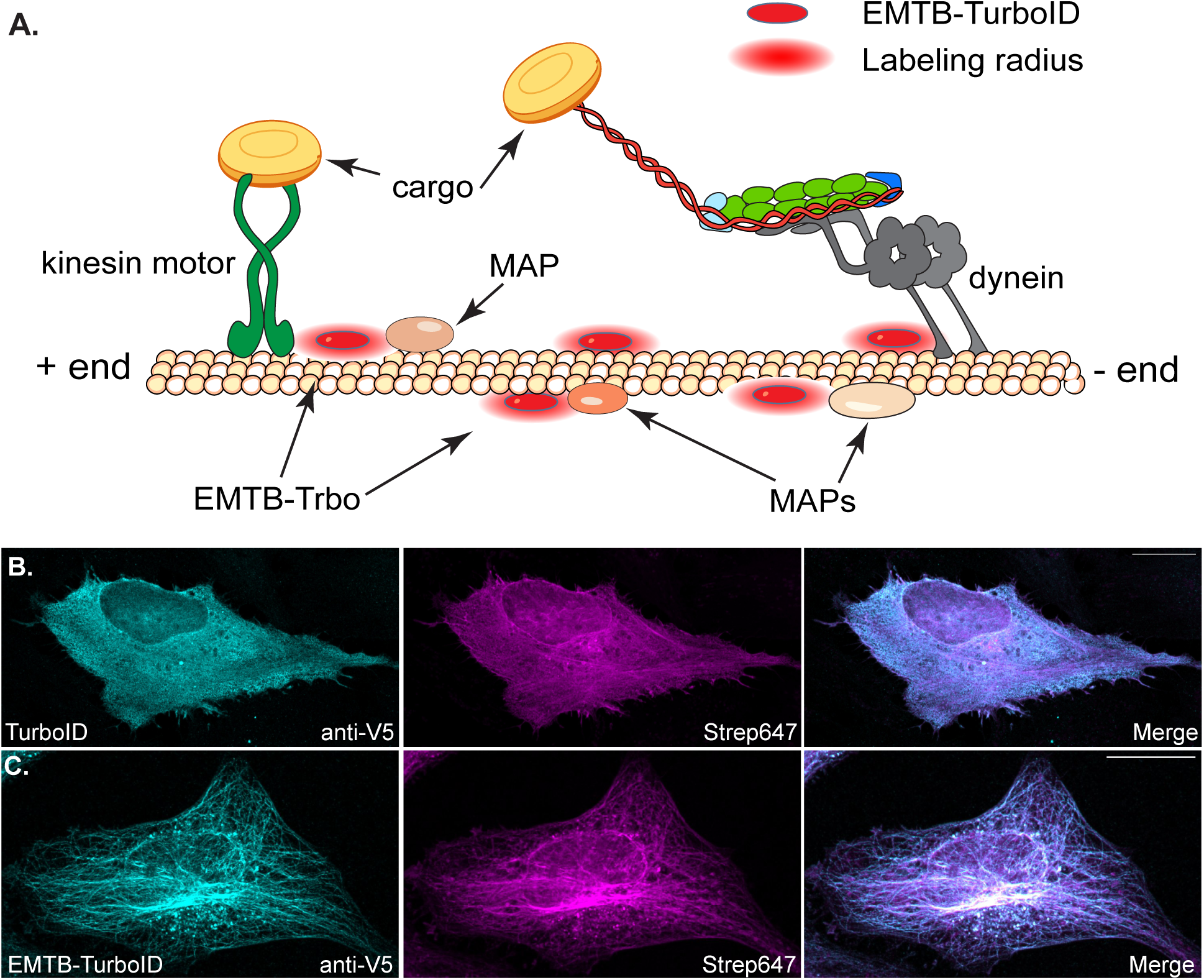
Validation of MT-ID. (A). A schematic illustrating the MT-ID approach. EMTB tagged with TurboID will bind to microtubules and result in the biotinylation of proximal microtubule associated proteins. (B-C). HeLa cells were transfected with a plasmid expressing either TubroID alone (B) or EMTB-TurboID (C). The cells were fixed and processed for immunofluorescence using a V5 antibody to detect TurboID (cyan). The cells were also probed using Streptavidin conjugated with Alexa647 to detect biotinylated proteins (magenta). A merged image is also shown. The scale bar is 20 microns.

We next generated stable cells containing either TurboID alone or EMTB-TurboID. HEK293 FLP In T-Rex cells were chosen for this experiment because this enabled us to target the transgenic constructs to a precise genomic locus and to specifically induce the expression of the construct using Tetracycline. In addition, we have successfully used this cell line in comparative proteomics experiments in the past (29). The cells were grown to scale and labeled with biotin. Next, biotinylated proteins were purified using streptavidin magnetic beads, and the bound proteins analyzed using mass spectrometry. The entire experiment was performed in triplicate. Proteins that displayed a 2-fold or greater enrichment with EMTB-Trbo versus the TurboID control and with a p value of at least 0.05 were considered microtubule interacting proteins (Fig. 2A, Supplemental Fig. 2A).

**Figure 2:**
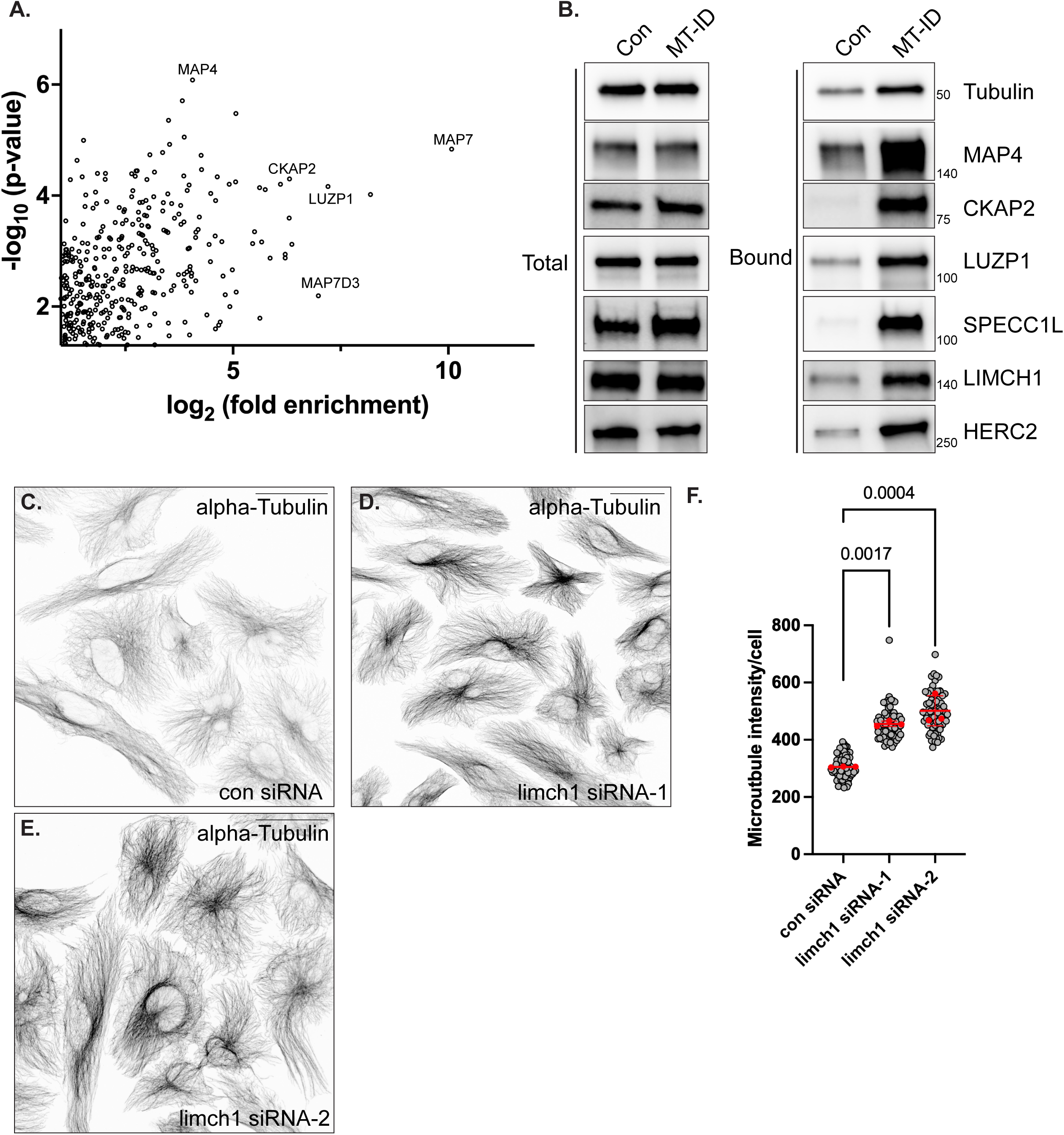
Identification of the microtubule interactome using MT-ID. (A) A volcano plot showing the candidates enriched at least two-fold with MT-ID versus the control and with a p value of at least 0.05. The full volcano plot is shown in Supplemental Fig. 2A. HEK293 FLP In T-Rex cells were used for this experiment. (B) Stable HEK293 FLP In T-Rex cells expressing either TurboID alone as the control or MT-ID were biotin labeled and harvested. Lysates were prepared from these cells and biotinylated proteins were purified. The bound proteins were analyzed by western blotting using the indicated antibodies. (C-F) HeLa cells were transfected with either a control siRNA (C) or siRNAs targeting LIMCH1 (D, E) The cells were fixed and processed for immunofluorescence using an antibody against alpha-Tubulin. Depletion of LIMCH1 results in higher levels of alpha tubulin staining per cell. The scale bar is 50 microns. (F) Quantification of the phenotype shown in panels C-E. Each red dot indicates the mean of a single biological replicate with the standard deviation for the three biological replicates indicated. The grey dots indicate the individual technical replicates. A one-way Anova was used for this analysis. The individual p values are indicated on the graph.

The most abundant classical or structural MAPs in our HEK293 cell MT-ID interactome were MAP7 isoforms (MAP7, MAP7D1, MAP7D2 and MAP7D3) and MAP4. The doublecortin-like protein, DCLK1, as well as MAP1A were also enriched with MT-ID in comparison to the control. In addition, PRC1 (Protein regulator of cytokinesis 1), a protein known to have a role in anti-parallel bundling of microtubules (30) was present in the MT-ID interactome. Surprisingly, although alpha and beta tubulin were recovered as specific interacting partners, their enrichment was not as high as that for MAP7 and MAP4. In order to validate these results, we repeated the experiment and analyzed the purified proteins using western blotting. Consistent with the proteomics result, MAP4 was highly enriched in the MT-ID sample in comparison to the control, with the enrichment of alpha-tubulin being more modest (Fig. 2B). Thus, although EMTB-TurboID binds to microtubules, the three-dimensional conformation of the complex likely facilitates better labeling of microtubule associated proteins in comparison to the microtubule subunits themselves.

Another class of proteins that were highly enriched in our microtubule interactome were centrosome and spindle associated proteins. We validated the specific enrichment of CKAP2 (Cytoskeleton-associated protein 2), a known spindle associated protein (31), with EMTB-TurboID. (Fig. 2B). As expected, CKAP2 was highly expressed in cells that were undergoing mitosis and localized to the mitotic spindle in these cells (Supplemental Fig. 2B). In addition, CKAP2 was also expressed at lower levels in interphase cells, wherein the protein was localized to microtubules (Supplemental Fig. 2C).

The microtubule and actin networks are highly interconnected, and several proteins are known to bind to both cytoskeletal filaments (32). Indeed, numerous proteins with established roles in actin-microtubule crosstalk such as Microtubule-actin cross-linking factor 1 (MACF1) and Dystonin (DST, also known as MACF2) (33) were highly enriched in our MT-ID interactome. We validated two of our top hits in this category, Leucine zipper protein1 (LUZP1) and Cytospin-A (SPECC1L) (Fig. 2B) (34, 35).

We next wished to determine whether the MT-ID approach could identify proteins not previously known to have microtubule-related functions. For this analysis, we chose to validate two proteins, HERC2 (E3 ubiquitin-protein ligase HERC2) and LIMCH1 (LIM and calponin homology domains-containing protein 1). Both proteins were enriched in the MT-ID interactome in comparison to the control. In addition, previous proteomic studies have shown that HERC2 was capable of immunoprecipitating tubulin isoforms, and that a subset of the protein localizes to centrosomes and centriolar satellites (36, 37). Although LIMCH1 has been shown to associate with Nonmuscle Myosin IIA (38), there are no reports indicating that it is also a microtubule associated protein. In order to validate the interaction of LIMCH1 and HERC2 with microtubules, we used an additional construct to confirm their microtubule association. The microtubule binding domain of Tau (MAPT), referred to as TMTB, fused to fluorescent proteins has also been shown to be useful in studying the dynamics of microtubules and to function in a comparable manner to EMTB (39). We therefore generated a stable cell line in which TMTB was fused to TurboID (TMTB-TurboID). In comparison to the control, MAP4, LUZP1, and CKAP2 were specifically enriched with both EMTB-TurboID and TMTB-TurboID (Supplemental Fig. 2D). The same pattern was observed for HERC2 and LIMCH1 (Supplemental Fig. 2D), thus confirming their association with microtubules.

We next determined the consequence of depleting LIMCH1 and HERC2. siRNAs were capable of depleting the target protein with no apparent effect on the level of total alpha tubulin by western blot (Supplemental Fig. 2E, F). Upon immunofluorescence analysis, we observed no dramatic change in the overall level and organization of microtubules in HeLa cells that were depleted of HERC2 (Supplemental Fig. 2G, GH). By contrast, HeLa cells depleted of LIMCH1 displayed higher levels of microtubule staining. This phenotype was observed using two independent siRNAs targeting LIMCH1 (Fig. 2C-F). Thus, LIMCH1 is a microtubule associated protein that regulates the steady state level of microtubules. The mechanism by which LIMCH1 performs this function is unknown and will be the focus of future studies.

### Comparison of the MT-ID interactome to the microtubule plus-end interactome

Microtubules are dynamic and polarized structures. Typically, microtubule growth occurs at the plus-end (40). By contrast, the microtubule minus-end is more stable and is often anchored within microtubule organizing centers present at the centrosome (6). In order to determine the spatial organization of the MT-ID interactome, we determined the interactome of the plus-end binding protein Eb3 (MAPRE3). Stable HEK FLP In T-Rex cells expressing either mScarlet3-TurboID or Eb3-TurboID were generated. The cells were grown to scale, labeled with biotin, and harvested. Biotinylated proteins were purified and identified using mass spectrometry as described in the previous section. As before, the entire experiment was performed in triplicate, and the same statistical parameters were used to determine Eb3 interacting proteins.

A significant fraction of the Eb3-TurboID interactome were either known Eb-interacting proteins, plus-end localized proteins, or proteins containing the SxIP motif typically found in plus-end localized proteins (40, 41). There was minimal overlap between the Eb3-TurboID interactome and the MT-ID interactome, with only 11% of proteins being shared between the two groups. In order to validate the Eb3 interactome, we repeated the experiment and probed the pellet using an antibody against CLIP1 (also known as CLIP170), the second most abundant protein in the Eb3 interactome. Consistent with the proteomics results, CLIP1 was abundantly detected in the Eb3-TurboID pellet compared to the control (Fig. 3B). In addition, a higher level of CLIP1 enrichment was observed with Eb3-TurboID versus MT-ID (Fig. 3B). The same pattern was observed for DCTN1, a component of the dynactin complex (Fig. 3A, B). In addition to functioning as an activator for the minus end motor, dynein, DCTN1 has also been shown to localize to the plus-end of microtubules and is thought to load dynein onto this region of the microtubule for minus-end transport (Fig. 3C) (42, 43). The known plus-end localized protein, TRIO (Triple functional domain protein) was also specifically enriched in the Eb3-TurboID pellet in comparison to the control and the MT-ID sample (Fig. 3B) (44). By contrast, the structural MAPs, MAP4 and MAP7 were more highly enriched in the MT-ID pellet (Fig. 3B). Based on these results, we conclude that most proteins identified in the MT-ID interactome correspond to microtubule lattice associated proteins.

**Figure 3:**
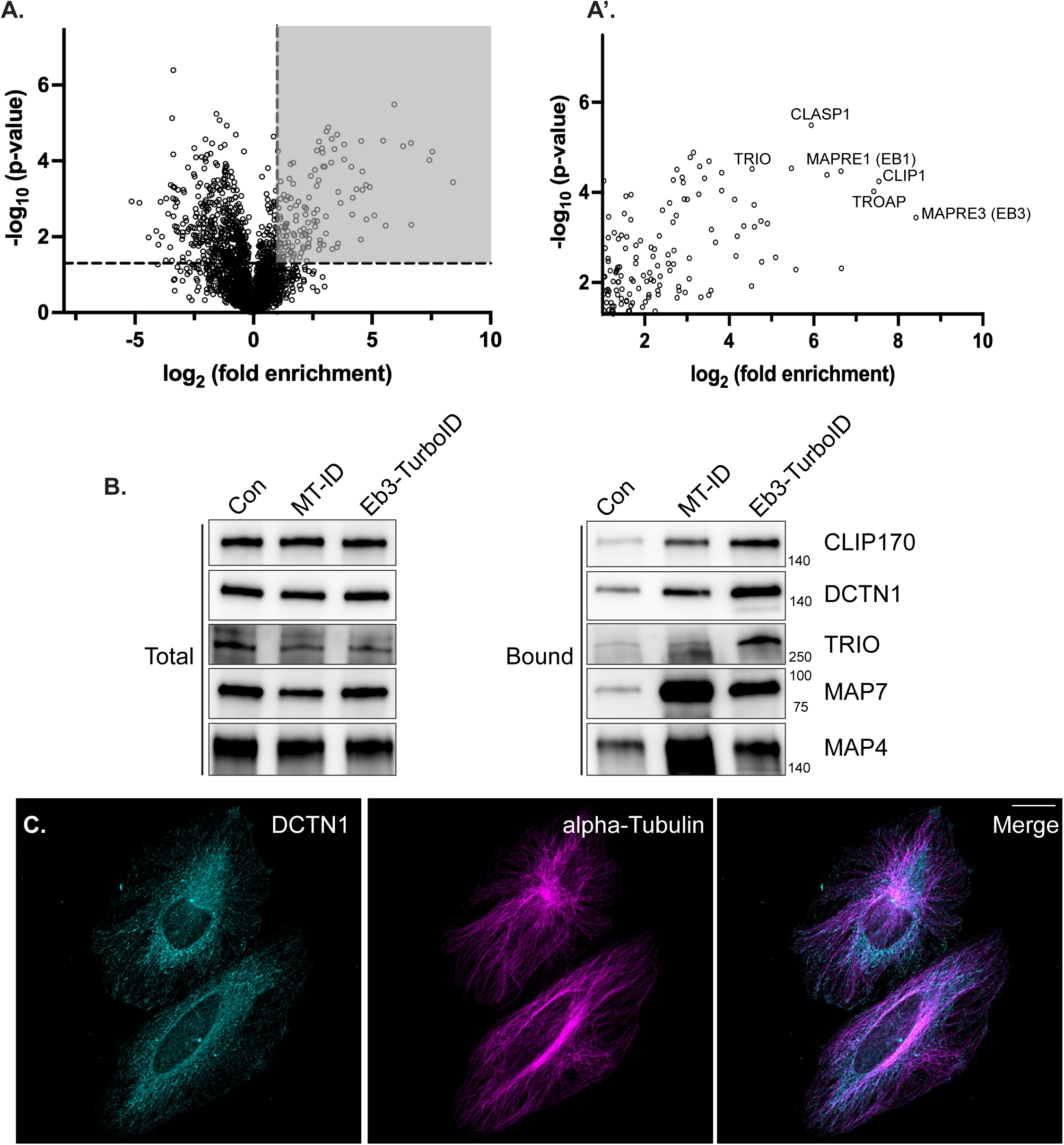
Comparison of the Eb3 and MT-ID interactomes. (A) A volcano plot showing proteins identified with Eb3-TurboID versus the mScarlet3-TurboID control. Proteins displaying at least two-fold enrichment with Eb3-TurboID and with a p value of at least 0.05 are indicated in A’. (B) Stable HEK293 FLP In T-Rex cells expressing either mScarlet3-TurboID, MT-ID (EMTB-TurboID) or Eb3-TurboID were biotin labeled and harvested. Lysates were prepared and biotinylated proteins were purified. The bound proteins were analyzed by western blotting using the indicated antibodies. Whereas CLIP170, DCTN1, and TRIO showed greater enrichment with Eb3-TurboID, MAP7 and MAP4 were more highly enriched with MT-ID. (C) HeLa cells were fixed and processed for immunofluorescence using antibodies against DCTN1 (cyan) and alpha-Tubulin (magenta). A merged image is also shown. The scale bar is 20 microns. DCTN1 localizes to the plus end of microtubules.

### MT-ID is capable of identifying the microtubule interactome in tissues

In the previous section, we determined the microtubule interactome in a cell line. In order to validate the general applicability of this approach, we wished to determine whether MT-ID could also be used to identify the microtubule interactome in tissues. The *Drosophila* egg chamber is an excellent model for studying microtubule organization and motor-based cargo transport (45). The egg chamber consists of 15 nurse cells and a single oocyte. Specification of the oocyte requires the polarized distribution of microtubules and cargo transport from the nurse cells into the oocyte is highly reliant on the microtubule minus end motor, dynein (46, 47). EMTB tagged with RFP has been used to examine MT dynamics in the egg chamber with no obvious negative effect on microtubule organization or cargo transport (48), thus suggesting that the MT-ID approach should work in this tissue. We therefore generated flies capable of expressing EMTB-TurboID using the UAS-GAL4 system. Expression of this construct using the germline specific nanos-Gal4 driver resulted in the biotinylation of microtubule proximal proteins (Fig. 4A, B). Encouraged by these findings, we scaled up the experiment for proteomic analysis. Ovaries were dissected from flies expressing either GFP-TurboID or EMTB-TurboID in the germline. Lysates were prepared and biotinylated proteins were purified using streptavidin magnetic beads. As before, the entire experiment was performed in triplicate, and the purified proteins were analyzed using mass spectrometry. Proteins that displayed a twofold or greater enrichment with EMTB-TurboID vs the GFP-TurboID control and with a p value of at least 0.05 were considered germline specific microtubule interacting proteins (Fig. 4C, C’).

**Figure 4:**
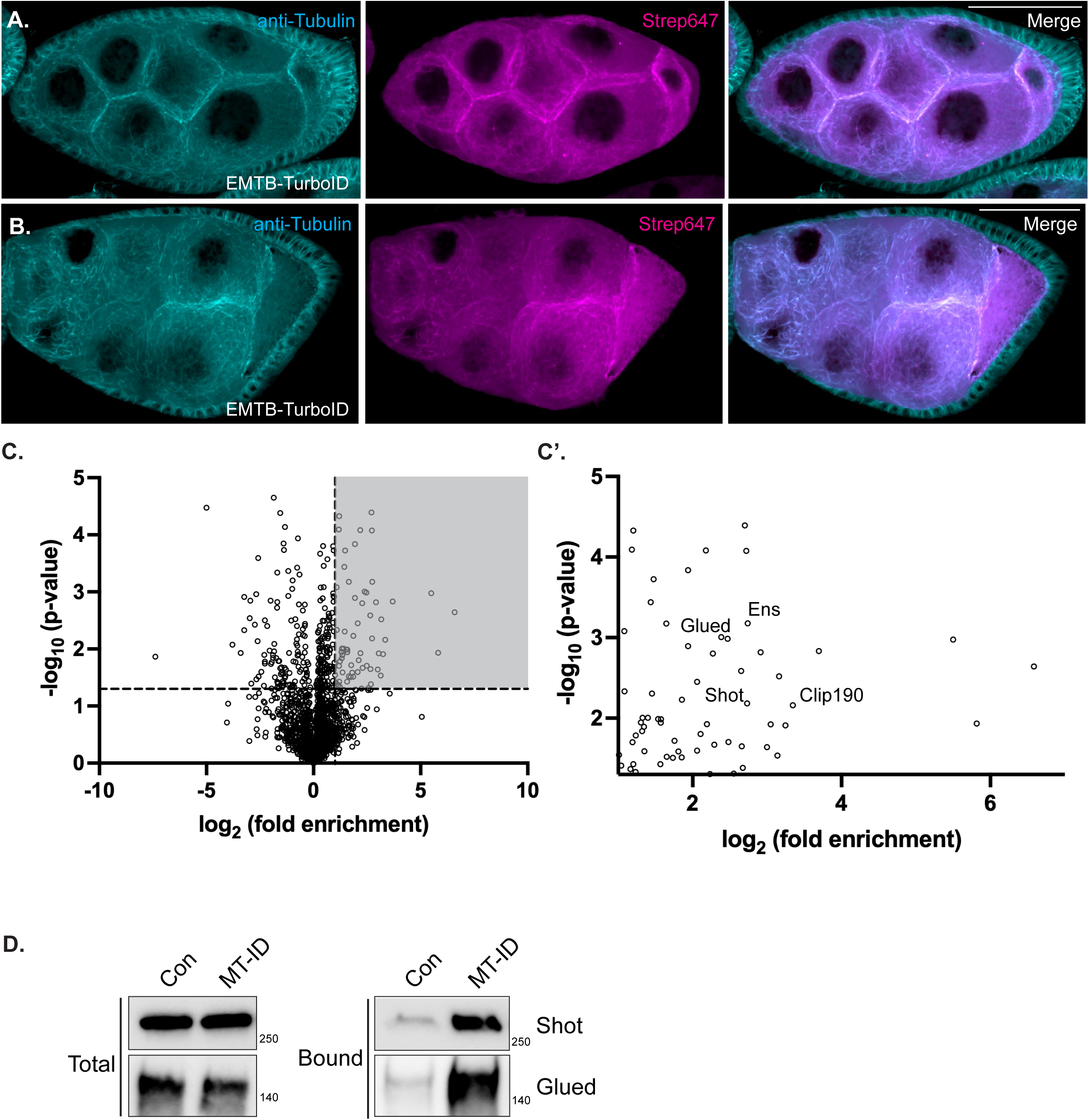
The MT-ID interactome in fly egg chambers. (A-B) Ovaries were dissected from flies expressing EMTB-TurboID using the nanos-Gal4 driver. The egg chambers were fixed and processed for immunofluorescence using an antibody against alpha-Tubulin (cyan). The egg chambers were also incubated with Streptavidin conjugated Alexa647 to detect biotinylated proteins (magenta). A merged image is also shown. The scale bar is 20 microns. (C) A volcano plot showing proteins identified with EMTB-TurboID versus the GFP-TurboID control. Proteins displaying at least two-fold enrichment with Eb3-TurboID and with a p value of at least 0.05 are shown in C’. (D) Ovaries were dissected from flies expressing EMTB-TurboID (MT-ID) or GFP-TurboID (con) in the germline. Lysates were prepared and biotinylated proteins were purified using streptavidin beads. Bound proteins were eluted and probed with the indicated antibodies. Consistent with the proteomics result, Shot and Glued were specifically enriched in the MT-ID sample.

Consistent with what was observed in HEK cells, the most abundant structural MAPs identified in our fly MT-ID interactome were Ensconsin and Map205, the fly orthologs of MAP7 and MAP4 respectively (49, 50). Another common component identified in both interactomes was the actin-microtubule crosslinking protein Shot (Fig. 4C’). The human homolog of Shot is Dystonin (DST), one of the most highly enriched proteins in the HEK MT-ID interactome. In order to validate this finding, we repeated the experiment and probed the purified proteins using an antibody against Shot. Consistent with the proteomics results, Shot was highly enriched in the ETMB-TurboID pellet in comparison to the control (Fig. 4D). There were also some notable differences between the fly and human MT-ID interactomes. For instance, the plus-end binding protein CLIP190 (CLIP1 in humans) was present in the fly MT-ID interactome but was not recovered in the HEK cell MT-ID interactome. In addition, the dynactin components, Glued and Dynamitin, (DCTN1 and DCTN2 in humans), were also enriched in the fly MT-ID interactome but were not significantly enriched in the HEK cell MT-ID interactome. We were able to validate this result using an antibody against Glued (Fig. 4D). Consistent with this finding, we have previously shown that Glued localizes to microtubules in the fly egg chamber (51), suggesting that it is indeed a bona fide microtubule associated protein in this tissue.

Collectively, these results indicate that the MT-ID approach can also be used to identify microtubule associated proteins in tissues. It should be noted, however, that the overall MT-ID interactome in the fly ovary was not as large as what was observed in HEK cells. The procedure of using fly ovaries for interactome analysis is challenging and further optimization (see discussion) will be needed to determine the full complement of microtubule associated proteins in this tissue.

### Development and validation of Act-ID

Much like the microtubule cytoskeleton, the actin cytoskeleton is also highly complex and regulated by a variety of binding proteins (14, 15). Current approaches are limited in their ability to globally define the actin interactome. To address this need, we developed Act-ID. Conceptually, Act-ID is similar to MT-ID. However, instead of a microtubule binding domain, the actin binding domain from the ITPKA protein (referred to as F-tractin) is fused to TurboID (Fig. 5A) (25). F-tractin fused to fluorescent proteins has been extensively used to study the dynamics of actin filaments in live cells (52–54). The recent structure of F-tractin revealed that a single mutation (F29A) reduces its ability to bind actin filaments (55). Consistent with these findings, F-tractin_wt_TurboID labels F-actin, whereas F-tractin_F29A_TurboID displays greatly reduced association with actin filaments (Fig. 5B, C).

**Figure 5:**
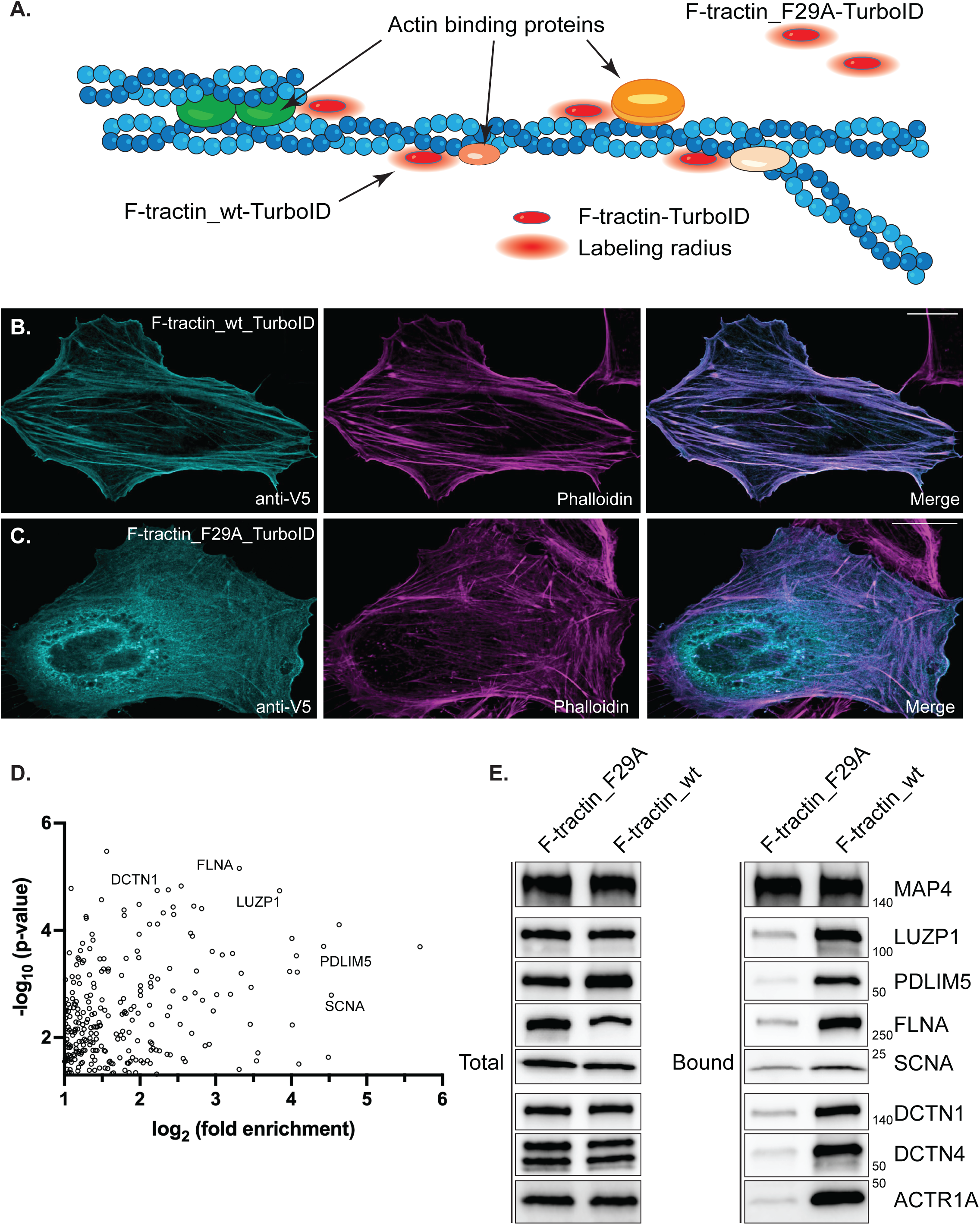
Validation of Act-ID. (A). A schematic illustrating the Act-ID approach. F-tractin_wt tagged with TurboID will bind F-actin and result in the biotinylation of proximal actin associated proteins. By contrast, F-tractin_F29A, also tagged in a similar manner, will not efficiently bind actin filaments and will serve as a control for the experiment. (B-C). HeLa cells were transfected with a plasmid expressing either TubroID-F-tractin_wt (B) or TubroID-F-tractin_F29A (C). The cells were fixed and processed for immunofluorescence using a V5 antibody to detect TurboID (cyan). The cells were also probed using Phalloidin conjugated with Alexa647 to detect actin filaments proteins (magenta). A merged image is also shown. The scale bar is 20 microns. (D) A volcano plot showing the candidates enriched at least two-fold with Act-ID versus the control and with a p value of at least 0.05. HEK293 FLP In T-Rex cells were used for this experiment. (E) Stable HEK293 FLP In T-Rex cells expressing either TubroID-F-tractin_wt (Act-ID) or TubroID-F-tractin_F29A were biotin labeled and harvested. Lysates were prepared from these cells and biotinylated proteins were purified. The bound proteins were analyzed by western blotting using the indicated antibodies.

Stable cells were generated using these constructs and the F-actin interactome was determined using biotin labeling, purification and mass spectrometry as described in previous sections. This analysis revealed a vast F-actin interactome with 261 proteins displaying a greater than two-fold interaction with F-tractin_wt in comparison to the F29A control (Fig. 5D). We validated the specific interaction of F-tractin_wt with a few top hits such as LUZP1, PDLIM5, FLNA, and SNCA (Fig. 5E), all of which have been previously shown to interact with actin (35, 56–58). By contrast, the abundant microtubule associate protein, MAP4, associated non-specifically with both WT and mutant F-tractin (Fig. 5E). Thus, Act-ID identifies genuine actin binding proteins.

Surprisingly, the Act-ID interactome contained several components of the Dynactin complex (DCTN1, DCTN2, DCTN4, DCTN5, ACTR1A) (Fig. 5E). As noted previously, the dynactin complex is a well-known regulator of the microtubule motor, dynein (59). Interestingly, the dynactin complex contains a central actin-like filament, which consists of eight copies of the actin related protein Arp1 (ACTR1A) and a single beta actin subunit (60–62). One possible explanation for why dynactin components were identified in the Act-ID interactome is because wild-type F-tractin-TurboID might be able to bind to the actin-like filament present within the dynactin complex. If true, this could result in the biotin labeling and purification of the dynactin complex. Another possibility is that because the dynactin complex is known to associate with microtubule plus-ends (42, 43), which are often found adjacent to the actin rich cell cortex, dynactin components are labeled by F-tractin at this site. Yet another possibility is that dynactin might actually be able to bind to traditional actin filaments. To distinguish between these possibilities, we depleted ACTR1A using siRNA. Two days after siRNA treatment, ACTR1A levels were greatly reduced (Supplemental Fig. 3A). However, at this time point, the level of DCTN1 and DCTN4 remained relatively unchanged (Supplemental Fig. 3A). This shorter time point of siRNA treatment was chosen because at longer time points, loss of one dynactin component destabilizes the rest of the complex (data not shown) (63). Depletion of ACTR1A using these conditions reduced the amount of DCTN1 and DCTN4 biotinylated and precipitated by wild-type F-tractin-TurboID (Supplemental fig. 3A). Based on this finding, the most likely conclusion is that F-tractin is able to associate with the actin-like filament found within dynactin. Further experiments will be needed to confirm whether F-tractin can bind to the central Arp1 filament in the dynactin complex.

We next wished to determine whether the Act-ID approach could identify proteins not previously known to have an actin related function. One of the most highly enriched candidates in our Act-ID interactome was the E3 ubiquitin ligase protein FBXO30 (F-box only protein 30). We chose FBXO30 for analysis because previous large scale affinity purification and proteomic studies have found this protein as a binding partner of beta actin (64–66). Indeed, consistent with these published findings and with our proteomic results, we were able to validate the association of FBXO30 with F-tractin-TurboID (Fig. 6A). In addition, GFP-FBXO30 was able to specifically co-precipitate beta actin from cell lysates (Fig. 6B). Because FBXO30 is an E3 ubiquitin ligase, we next determined whether depletion of this protein would affect the total level of beta actin. Although FBXO30 could be effectively depleted using two independent siRNAs, the total level of beta actin in cell lysates was unchanged (Supplemental Fig. 3B). We next tested whether loss of FBXO30 would result in changes to the organization of F-actin filaments. Interestingly, in contrast to cells transfected with a control siRNA, cells that were depleted of FBXO30 displayed reduced cortical actin fibers (Fig. 6C-E, arrow). Instead, these cells contained more central actin stress fibers (Fig. 6C-E, arrowhead). Consistent with this finding, cells that were depleted of FBXO30 had an increased number of focal adhesions, visualized using the maker protein Vinculin (Fig. 6C’-E’, F). Although depletion of FBXO30 resulted in increased focal adhesions, the total level of Vinculin was unaffected in these lysates (Supplemental Fig. 3B). Thus, FBXO30 does not appear to directly target either beta actin or Vinculin for proteasomal degradation. Collectively, our results suggest that FBXO30 is a novel actin binding protein that functions to restrain the formation of focal adhesions.

**Figure 6:**
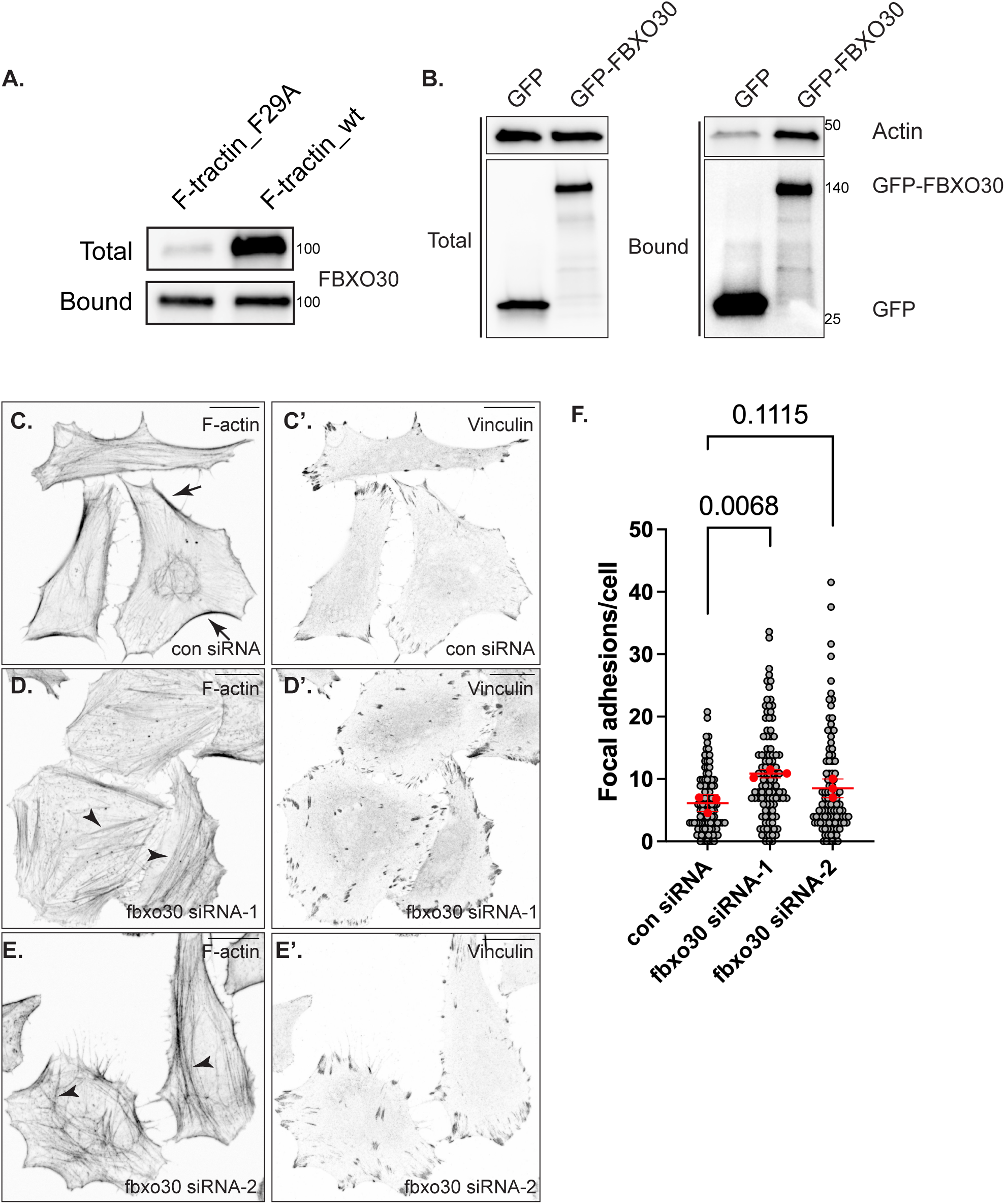
FBXO30 is a novel actin binding protein. (A). Stable HEK293 FLP In T-Rex cells expressing either TubroID-F-tractin_wt (Act-ID) or the control were biotin labeled and harvested. Lysates were prepared and biotinylated proteins were purified. The bound proteins were analyzed by western blotting using an antibody against FBXO30. (B). HEK293 FLP In T-Rex cells were transfected with a plasmid expressing either GFP alone or GFP-FBXO30. Two days after transfection, lysates were prepared and the GFP tagged protein was precipitated using GFP-Trap beads. After appropriate wash steps, the bounds proteins were analyzed by western blotting using antibodies against beta-actin and GFP. Beta-actin specifically co-precipitates with GFP-FBXO30. (C-F) HeLa cells were transfected with either a control siRNA (C) or siRNAs targeting FBXO30 (D, E) The cells were fixed and processed for immunofluorescence using an antibody against Vinculin (C’, D’, E’). The cells were also incubated with Phalloidin conjugated with Alexa647 to reveal the actin network (C, D, E). Arrows indicate cortical bundles of actin and arrow-heads indicate internal stress fibers. (F) Quantification of the focal adhesion phenotype shown in panels C’-E’. Each red dot indicates the mean of a single biological replicate with the standard deviation for the three biological replicates indicated. The grey dots indicate the individual technical replicates. A one-way Anova was used for this analysis and the individual p values are indicated.

## DISCUSSION

The proper functioning of the microtubule and actin cytoskeletal networks relies on a diverse set of proteins that bind these filaments to regulate their dynamics, their arrangement into higher-order structures, and their ability to respond to a variety of stimuli (2, 14). Our goal, therefore, was to devise an approach to identify these cytoskeletal-associated proteins in cells and tissues under physiological conditions, without the use of crosslinkers or reagents meant to artificially stabilize the network. To accomplish this task, we used an in vivo proximity biotin ligation approach in which the biotin ligase TurboID was fused to either a microtubule- or actin-binding domain. We refer to these strategies as MT-ID (for microtubules) and Act-ID (for actin).

MT-ID was capable of identifying a vast microtubule interactome. Structural MAPs such as MAP7 isoforms and MAP4 were highly enriched in the MT-ID interactome. We also validated the association of known microtubule associated proteins and proteins known to interact with both microtubule and actin networks. In addition, we provide evidence that LIMCH1 is a novel microtubule associated protein. Depletion of LIMCH1 results in greater microtubule density per cell. The mechanism by which LIMCH1 performs this function is an open question and will be a topic for future studies.

One protein that we expected to abundantly detect in the MT-ID interactome was the heavy chain of the plus-end motor kinesin-1, KIF5B. Although KIF5B could be detected in the MT-ID interactome, it was minimally enriched in comparison to the control. Unlike structural MAPs, motor proteins only transiently interact with the microtubule network during the process of cargo transport. We thus wondered whether KIF5B would display a greater enrichment with MT-ID if we prolonged its interaction with microtubules. This was done by incubating the cells with AMP-PNP, a non-hydrolyzeable ATP analog, which prevents kinesin-1 motors that are bound to microtubules from detaching (67, 68). Indeed, treatment with AMP-PNP greatly increased the enrichment of KIF5B in the MT-ID sample (Supplemental Fig. 2I). Thus, one limitation of MT-ID is its ability to efficiently identify proteins such as motors that only transiently interact with the microtubule network.

Although EMTB-TurboID efficiently localized to microtubules in interphase cells, it was less efficient at localizing to microtubules in cells that were undergoing mitosis (data not shown). This is consistent with the finding that MAP7 displays cell-cycle specific localization patterns (16). In addition, MAP7 is phosphorylated during mitosis, and this modification reduces its affinity for microtubules (3, 69). Thus, it is likely that microtubule associated proteins with a specific role in mitosis are under-represented in our MT-ID interactome. One way to circumvent this limitation is to generate a mutant version of EMTB in which the conserved phosphorylation sites are mutated. This should restore microtubule binding during mitosis and enable the identification of microtubule associated proteins with specific roles during this stage of the cell cycle.

In addition to identifying the microtubule interactome in a cells line, we were also able to utilize this approach in tissues. We chose to test the utility of MT-ID using the *Drosophila* egg chamber as a model. The fly ovary is a well-characterized tissue in which microtubule motor-based transport is critical for delivering cargos from the nurse cells into the oocyte (45, 47, 70). Although we were able to identify microtubule associated proteins such as the fly homologs of MAP7 and MAP4 (Ensconsin and Map205 respectively), the ovary MT-ID interactome contained far fewer candidate proteins than what was identified in HEK cells. Although it is possible that fewer proteins are associated with microtubules in this tissue, it is more likely that technical limitations contributed to this finding. In contrast to cells, which can be grown to scale and easily harvested for experiments, ovaries must be dissected from individual flies. In addition, we intentionally expressed the EMTB-TurboID construct at a low level in this tissue, because expression at a high level resulted in microtubule bundling (data not shown). Also, in keeping with our previous studies (71), we did not add exogenous biotin to the fly food. However, given the low expression level of the MT-ID construct in this tissue, exogenous addition of biotin will likely improve the recovery of microtubule associated proteins that we failed to identify in the current study.

Similar to the MT-ID approach, we also demonstrate that Act-ID was capable of identifying an extensive actin interactome. For this strategy, we chose to use F-tractin as our actin binding domain because a single point mutation in F-tractin has been shown to significantly reduce actin filament binding (55). Thus, this mutant served as an excellent control for binding and specificity. However, in theory, other actin binding domains such as Lifeact (72, 73) or Utrophin (74) fused to TurboID should also produce similar results. In addition to identifying known actin interacting proteins, the Act-ID interactome also contained several components of the dynactin complex. This was a surprising finding because dynactin is an activator of the dynein motor (10), and despite many years of study, has not been implicated as a classical actin binding protein. However, the dynactin complex consists of several subunits built around an actin-like filament composed of Arp1, Arp11, and a single beta actin subunit (60–63). Thus, the most likely explanation is that F-tractin binds to the actin-like filament within the dynactin complex, resulting in the biotinylation and purification of these components. Consistent with this notion, depletion of Arp1 (ACTR1A) reduces the level of DCTN1 and DCTN4 brought down by Act-ID.

One of the top candidates in the Act-ID interactome was the poorly characterized protein FBXO30. FBXO30 is an F-box protein and it functions as the substrate recognition component of the SCF (Skp1-Cullin-F-box) E3 ubiquitin ligase complex (75, 76). Although the specific substrates targeted by FBXO30 for proteasomal degradation are not known, FBXO30 is significantly upregulated under conditions of skeletal muscle atrophy and its high level of expression is thought to contribute to this muscle phenotype (77). We chose to perform follow-up studies on FBXO30 because several large-scale proteomic screens identified it as an interacting partner for beta actin (64–66). In line with these studies, we found that GFP-FBXO30 was able to co-precipitate beta actin from cell lysates. While depletion of FBXO30 did not affect the total level of actin, the organization of actin filaments were altered and the depleted cells displayed an increased number of focal adhesions. Future studies will seek to define the substrates of FBXO30 and the mechanism by which FBXO30 loss induces focal adhesion formation.

The goal of our study was to develop the tools to analyze the protein interaction landscape of the microtubule and actin cytoskeletal networks. The next step would be to use these tools to understand how the cytoskeletal interactome changes under different physiological conditions. For instance, osmotic changes have been shown to affect the post-translational modification (PTM) and MAP association profile of microtubules (78). These experiments were performed using antibodies against specific MAPs and microtubule PTMs (78). With MT-ID it should be possible to define global changes that occur to the microtubule interactome when cells are exposed to different osmotic conditions. In addition, it was recently demonstrated that oxidative stress results in the dynein-dependent re-localization of several organelles, yet the mechanism that resulted in dynein hyperactivity is not known (79). MT-ID can be used to determine whether oxidative stress results in changes to the microtubule interactome and whether these changes in turn contribute to dynein activation. Finally, mutations in certain cytoskeletal associated proteins, motors, or their accessory factors are associated with disease (80–82). MT-ID and Act-ID will be powerful tools to determine how these disease associated mutations affect the cytoskeletal interactome.

## ACKNOWLEDGEMENTS

We are grateful to Drs. Vladimir Gelfand and Wen Lu for the antibody against *Drosophila* Glued and for helpful discussion of this project. We are also grateful to Pritha Bagchi for her assistance with the mass-spectrometry analysis. This work was supported by a grant from the National Institutes of Health to G.B.G (NIGMS, R35GM145340). This work was supported in part by the Emory Integrated Proteomics Core (RRID:SCR_023530) and the Augusta University Cell Imaging Core (RRID:SCR_026799).

## MATERIALS AND METHODS

### DNA constructs

Synthesized DNA fragments were generated by Genewiz/Azenta. All final plasmids used in this work were sequenced by Plasmidsaurus. Plasmid sequences as well as detailed cloning strategies will be provided upon request. The following constructs were generated in the pCDNA-FRT-TO vector from Thermo Fisher: TurboID alone, EMTB-TurboID, TMTB-TurboID, Eb3-TurboID, mScarlet3-TurboID, TurboID-F-tractin_wt and TurboID-F-tractin_F29A. In addition, for studies in *Drosophila* tissues, EMTB-TurboID and GFP-TurboID were cloned into the pUASp-attB vector.

### Cell culture

HeLa, Cos7, U2OS and hTERT-RPE cells were authenticated and obtained from ATCC. HeLa, Cos7 and USOS cells were cultured in Dulbecco’s Modified Eagle Medium (DMEM) supplemented with 10% fetal bovine serum and 1% penicillin/streptomycin (Thermo Fisher Scientific). hTERT-RPE cells were cultured in DMEM/F-12 50/50 supplemented with 10% fetal bovine serum and 1% penicillin/streptomycin (Thermo Fisher Scientific).

HEK Flp-In T-REx 293 cells were obtained from Thermo Fisher Scientific and were cultured according to the instructions provided by the manufacturer. Stable cells were generated by culturing the transfected HEK Flp-In T-REx 293 in 100ug/ml Hygromycin B (Gibco) for 14 days, with fresh media changes every 3 days. Expression of the all constructs except Eb3-TurboID were induced using 1ug/ml Tetracyline for 24 hours. For Eb3-TurboID, expression was induced with the same concentration of Tetracycline but for only 6 hours. This was done to reduce expression level and prevent the spreading of Eb3-TurboID away from the microtubule plus end.

### Fly stocks

The pUASt-EMTB-TurboID and GFP-TurboID constructs were inserted at the ZH-68E site (Bloomington stock center; #24485, donor Konrad Basler & Johannes Bischof). The strains were injected by BestGene Inc. Expression of the pUASt-EMTB-TurboID construct was driven using the following driver: P{w[+mC]=GAL4::VP16-nos.UTR}CG6325[MVD1] (Bloomington Stock center, #4937; donor Ruth Lehmann). The GFP-TurboID control strain was generated by cloning the coding sequences of GFP and TurboID into a plasmid with a maternal promoter vector and 3’UTR. Fly stocks and crosses used for these experiments were maintained at 25^0^C.

### Antibodies and Reagents

The following antibodies were used in this study:

V5 (Thermo Fisher; R960-25, 1:1000 for immunofluorescence, 1:10,000 for western), alpha tubulin (Millipore Sigma, 10-013-CM, 1:40,000 for western, 1:4000 for immunofluorescence), MAP4 (Proteintech, 11229-1-AP; 1:1500 for westerns), CKAP2 (Proteintech, 25486-1-AP; 1:4000 for westerns), LUZP1 (Proteintech, 17483-1-AP; 1:5000 for western), SPECC1L (Proteintech, 25390-1-AP, 1:600 for western), LIMCH1 (Novus biologicals, NBP1-32614, 1:3000 for western), HERC2 (Proteintech, 27459-1-AP, 1:500 for western), DCTN1 (Thermo Fisher, PA5-21289, 1:250 dilution for immunofluorescence, 1:2500 dilution for western), CLIP1 (Proteintech, 23839-1-AP; 1:3000 for western), TRIO (Proteintech, 31192-1-AP, 1:500 for western), Shot (Developmental studies hybridoma bank, anti-Shot mAbRod1), Glued (1:5000 for western, gift from V. Gelfand), PDLIM5 (Proteintech, 10530-1-AP, 1:1000 for western), FLNA (Proteintech, 67133-1-Ig, 1:10,000 for western), SCNA (Proteintech, 10842-1-AP, 1:800 for western), DCTN4 (Fisher 50-156-9205), ACTR1A (Proteintech, 11023-1-AP, 1:1000 for western), FBOX30 (Proteintech, 28039-1-AP, 1:5000 for western), beta actin (Proteintech, 20536-1-AP, 1:5000 for western), Vinculin (Proteintech, 26520-1-AP; 1:4000 for immunofluorescence, 1:40,0000 for western), KIF5B (Proteintech, 21632-1-AP, 1:5000 for western), GFP (Proteintech, pabg1, 1:1000 for western). CALD1 (Proteintech, 66693-1-Ig, 1:5000 for western), goat anti-mouse Alexa 488 plus (Thermo Fisher, A32723TR, 1:400), goat anti-mouse Alexa 555 plus (Thermo Fisher, A32727, 1:400), goat anti-rabbit Alexa 488 plus (Thermo Fisher, A32731TR, 1:400), and goat anti-rabbit Alexa 555 plus (Thermo Fisher, A32732, 1:400). F-actin was visualized using Phalloidin-Alexa647 (Thermo Fisher, A22287, 1:600) and biotinylated proteins were visualized using Streptavidin Alexa647 (Thermo Fisher, S21374, 1:1200).

All siRNAs used were obtained from Thermo Fisher. The following siRNAs were used in this study: non targeting control (catalog # 4390843); herc2 siRNA (4392420, assay ID s17064); limch1 siRNA-1 (4427038, assay ID s22806); limch1 siRNA-2 (4392420, assay ID s22805), fbxo30 siRNA-1 (4392420, assay ID s38493); fbxo siRNA-2 (4427037, assay ID s38492) and actr1A siRNA (4427037, assay ID s19692).

GFP trap beads (Proteintech, catalog # gta) were used for the co-immunoprecipitation experiments.

### DNA and siRNA transfections

The Qiagen Effectene reagent was used for transfecting DNA into HeLa, HEK Flp-In T-REx 293, and Cos7 cells. Lipofectamine LTX plus reagent from Thermo Fisher was used for transfecting DNA into U2OS cells and hTERT-RPE cells. siRNA was transfected into cells using RNAi Max from Thermo Fisher. In all cases, manufacturer instructions were followed. For transient expression, cells were fixed and processed for immunofluorescence one day following transfection. For siRNA transfection, the cells were processed either 2 days or 3 days after transfection.

### Purification and analysis of biotinylated proteins

For small-scale assays, HEK Flp-In T-REx 293 cells were induced to express target constructs using tetracycline (1 μg/ml, 24 hours) and subsequently exposed to biotin (500 μM) for 10 minutes. Cells were collected and lysed by resuspension in RIPA buffer (50 mM Tris-Cl [pH 7.5], 150 mM NaCl, 1% NP-40, 1 mM EDTA) supplemented with Halt Protease inhibitor cocktail (Thermo Fisher Scientific). Binding reactions utilized 1 mg total protein incubated with 15 μl High-Capacity Streptavidin Agarose beads (Thermo Fisher Scientific) in RIPA buffer overnight at 4°C. Following four washes with RIPA buffer, bound proteins were released in Laemmli buffer and subjected to western blot analysis. Western blot imaging was performed using a Bio-Rad ChemiDoc MP system.

For proteomic analyses, identical cells were cultured in 10 cm dishes, and 5 mg total protein was employed per purification. Biotinylated proteins were captured using 70 μl Streptavidin magnetic beads (Thermo Fisher Scientific) during overnight incubation at 4°C. Samples underwent extensive washing with 1 ml volumes as follows: three washes with RIPA buffer, three washes with high-salt RIPA buffer (50 mM Tris-Cl [pH 7.5], 1 M NaCl, 1% NP-40, 1 mM EDTA), three additional RIPA buffer washes, and four PBS washes. All experiments were performed in triplicate. Following final washes, beads were resuspended in 70 μl PBS and transported on dry ice to the Emory Integrated Proteomics Core.

### Mass spectrometry

The mass spectrometry was performed at the Emory Integrated Proteomics Core (RRID:SCR_023530).

#### On-bead digestion

The on-bead protein digestion procedure was conducted following an established protocol (83). Beads were treated with digestion buffer (50 mM NH4HCO3), then subjected to reduction with 1 mM dithiothreitol (DTT) for 30 minutes at ambient temperature, after which 5 mM iodoacetamide (IAA) was introduced. The sample underwent alkylation for 30 minutes at ambient temperature while protected from light. Enzymatic digestion was performed sequentially using 1 µg lysyl endopeptidase (Wako) overnight at ambient temperature, followed by overnight digestion with 1 µg trypsin (Promega) under the same conditions. The generated peptides underwent desalting with an HLB column (Waters) before vacuum drying.

#### LC-MS/MS

Liquid chromatography-tandem mass spectrometry data collection followed a previously described method (84). Peptides were reconstituted in loading buffer (0.1% trifluoroacetic acid, TFA) and chromatographically separated using a Water’s Charged Surface Hybrid (CSH) column (internal diameter: 150 µm; length: 15 cm; particle size: 1.7 µm). Chromatographic separation was achieved on an EVOSEP system utilizing the 15 samples per day preset gradient program, with detection performed on a Q-Exactive Plus Hybrid Quadrupole-Orbitrap Mass Spectrometer (ThermoFisher Scientific). Each acquisition cycle consisted of one full MS scan and 20 subsequent data-dependent MS/MS scans. Full MS scans were acquired in profile mode at 70,000 resolution (at m/z 200) across 400-1600 m/z with an AGC target of 3 × 10⁶ and 100 ms maximum ion time. HCD fragmentation spectra were obtained at 17,500 resolution (at m/z 200) with parameters including 1.6 m/z isolation width, 28% collision energy, 1 × 10⁵ AGC target, and 100 ms maximum ion time. Previously sequenced precursors were dynamically excluded for 30 seconds. Ions with charge states of +1, +7, +8, or higher were excluded from analysis.

#### MaxQuant

The label-free quantification workflow was based on a previously published method (Seyfried, Dammer et al. 2017). Database searching was conducted using Andromeda (integrated within MaxQuant) against the 2022 human UniProtKB/Swiss-Prot database containing 20,387 target sequences. Variable modifications included methionine oxidation (+15.9949 Da), asparagine/glutamine deamidation (+0.9840 Da), and protein N-terminal acetylation (+42.0106 Da), with a maximum of 5 modifications per peptide; carbamidomethylation of cysteine (+57.0215 Da) was set as a fixed modification. Search parameters required fully tryptic peptides with a maximum of 2 missed cleavages permitted. Mass tolerance was set at ±20 ppm before calibration and ±4.5 ppm following internal calibration. Additional parameters included maximum peptide mass of 6,000 Da, minimum peptide length of 6 residues, and MS/MS mass tolerances of 0.05 Da (orbitrap) and 0.6 Da (ion trap). A 1% false discovery rate threshold was applied for peptide-spectrum matches, proteins, and site identifications.

#### For quantification

second-pass re-quantification was enabled following initial protein identification; cross-run MS1 feature matching was activated with a 0.7-minute retention time tolerance window following alignment over a 20-minute search interval. Protein abundance was determined from summed peptide intensities calculated by MaxQuant, utilizing only unique and razor peptides for protein-level quantification.

### Co-Immunoprecipitation

HEK293 FLP-in T-Rex cells were transfected with plasmids expressing either GFP alone or GFP-FBXO30. 1mg of total protein was used in the binding experiment with 15ul of GFP-Trap agarose beads (ChromoTek, ProteinTech) in binding buffer (50mM Tris pH 7.5, 150mM NaCl, 0.2mM EDTA, and 0.05% NP40). The binding was performed at 4^0^C for 2 hours. The samples were then washed four times using binding buffer, bound proteins were eluted in Laemmli buffer and analyzed by western blotting using the indicated antibodies.

### Immunofluorescence

Cells (HeLa, Cos7, U2OS, and hTERT-RPE) adhered to glass coverslips were fixed using either 4% formaldehyde for 5 minutes at room temperature (Figures 5, 6) or with methanol at -20^0^C for 10 minutes (Figures 1, 2, 3, Supplemental figure 1 and Supplemental figure 2). For the experiment in figure 4, egg chambers were first incubated in a PIPES extraction buffer as described previously (83). After this incubation, the tissues were fixed using 4% formaldehyde for 15 minutes at room temperature. After fixation, cells and tissues were permeabilized by incubating with PBST (PBS containing 0.1% TritonX-100) for 5 minutes. Next the samples were blocked for 1 hour at room temperature using 5% normal goat serum (Thermo Fisher). Cells were incubated overnight at 4^0^C with primary antibody in blocking solution. The next day, the coverslips were washed three times with PBS. The cells were then incubated with secondary antibody diluted in blocking solution for 1 hour at room temperature. Following this, the cells were washed three times with PBS, stained with DAPI (Thermo Fisher), and mounted onto slides using Prolong Diamond antifade reagent (Thermo Fisher). In figures 5 and 6 the cells were also counterstained with Phalloidin Alexa647 to reveal the actin network.

### Microscopy

All imaging experiments were performed at the Augusta University Cell Imaging Core (RRID:SCR_026799). Fixed images were captured on either an inverted Leica Stellaris confocal microscope or an inverted Nikon AXR confocal microscope equipped with the NSPARC detector. Images were processed for presentation using Fiji, Adobe Photoshop, and Adobe Illustrator.

### Quantifications

Microtubule intensity per cell was calculated using Fiji/ImageJ. This was done by using the threshold feature to define regions of interest (ROI). The average signal intensity for the alpha-tubulin channel was then measured in each of these ROIs. The number of focal adhesions per cells was measured using the G3 analysis pipeline of Nikon elements. The DAPI signal for cell nuclei as well as the F-actin signal was used to create a mask for each cell. The number of focal adhesions per cell were then counted using the spots feature.

---

**Supplemental figure 1.**

(A-C. Cos7 (A), U2OS (B) or hTERT-RPE (C) cells were transfected with a plasmid expressing either EMTB-TurboID. The cells were fixed and processed for immunofluorescence using a V5 antibody to detect TurboID (cyan). The cells were also probed using Streptavidin conjugated with Alexa647 to detect biotinylated proteins (magenta). A merged image is also shown. The scale bar is 20 microns.

**Supplemental figure 2.**

(A) A volcano plot showing proteins identified with MT-ID versus the control. Proteins displaying at least two-fold enrichment with MT-ID and with a p value of at least 0.05 are shaded in grey.

(B-C) HeLa cells were fixed and processed for immunofluorescence using antibodies against CKAP2 (cyan) and alpha-Tubulin (magenta). A merged image is also shown. The scale bar is 20 microns.

(D) Stable HEK293 FLP In T-Rex cells expressing either TurboID alone, EMTB-TurboID or TMTB-TurboID were biotin labeled and harvested. Lysates were prepared from these cells and biotinylated proteins were purified. The bound proteins were analyzed by western blotting using the indicated antibodies.

(E-F) HEK293 FLP In T-Rex cells were transfected with a control siRNA or siRNAs targeting HERC2 (E) or LIMCH1 (F). Three days after transfection, cells were harvested and the lysates were analyzed using the indicated antibodies.

(G-H) HeLa cells were transfected with a control siRNA (G) or siRNAs targeting HERC2 (H). Three days after transfection, cells were fixed and processed for immunofluorescence using an antibody against alpha-Tubulin. Microtubule levels are similar between control and HERC2 depleted cells.

(I) Stable HEK293 FLP In T-Rex cells expressing either TurboID alone or EMTB-TurboID were incubated with vehicle (DMSO) or with AMP-PNP, a non-hydolysable ATP analog. The cells were biotin labeled and harvested. Lysates were prepared, biotinylated proteins were purified and analyzed by western blotting using an antibody against KIF5B. EMTB-TurboID is able to more efficiently biotinylate KIF5B when the cells are treated with AMP-PNP.

**Supplemental figure 3.**

(A) Stable HEK293 FLP In T-Rex cells expressing TubroID-F-tractin_wt (Act-ID) were transfected with either a control siRNA or an siRNA targeting ACTR1A. Two days after transfection, the cells were labeled with biotin had harvested. Lysates were prepared, biotinylated proteins were purified and analyzed by western blotting using the indicated antibodies.

(B) HEK293 FLP In T-Rex cells were transfected with a control siRNA or siRNAs targeting FBXO30. Three days after transfection, cells were harvested and the lysates were analyzed using the indicated antibodies.

## Notes

### Competing Interest Statement

The authors have declared no competing interest.

