## Supplementary figures and images for "Proteomic profiling of cytoskeletal interactomes using MT-ID and Act-ID"

### Supplemental Figure1

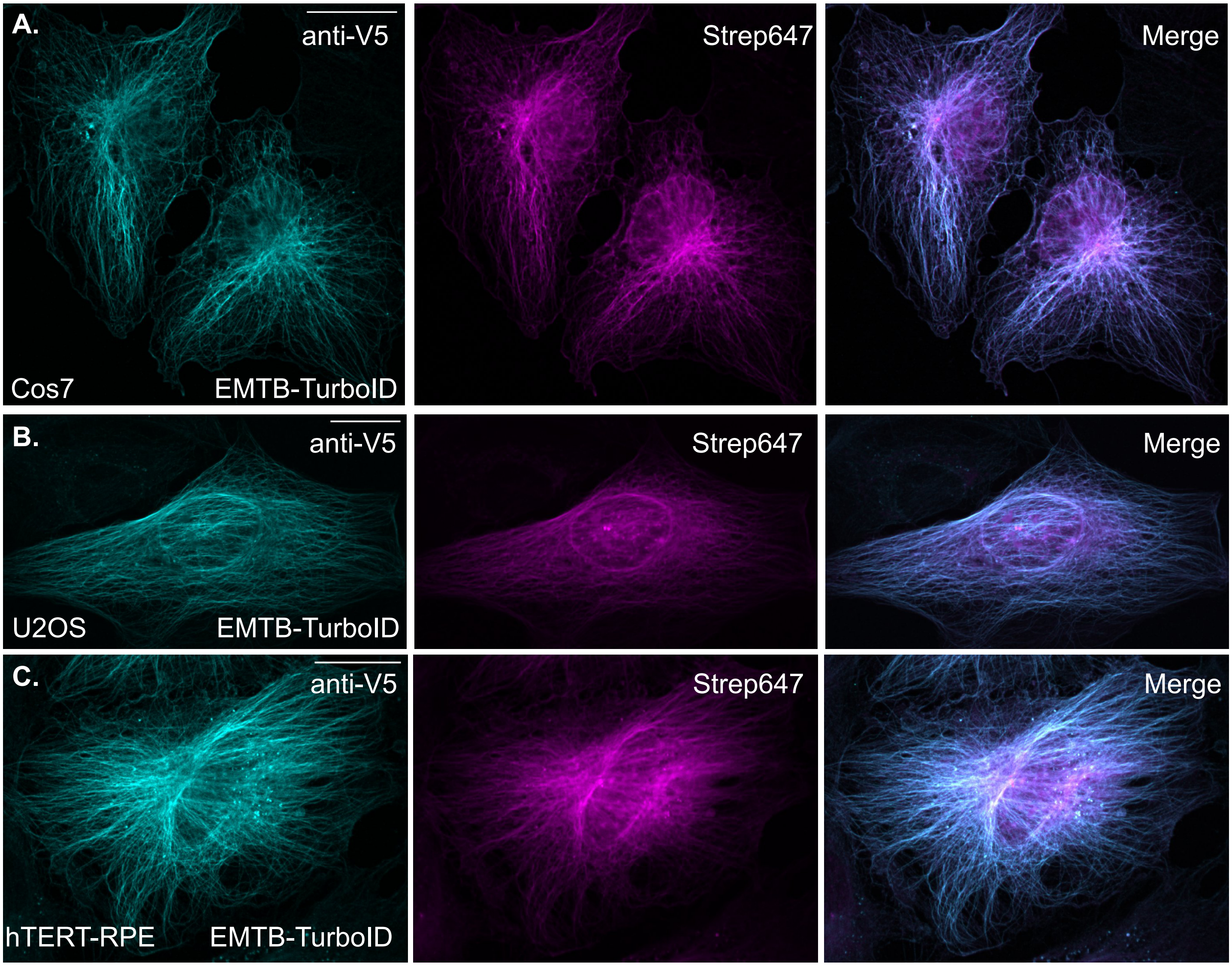


Supplemental Fig. 1

### Supplemental Figure2

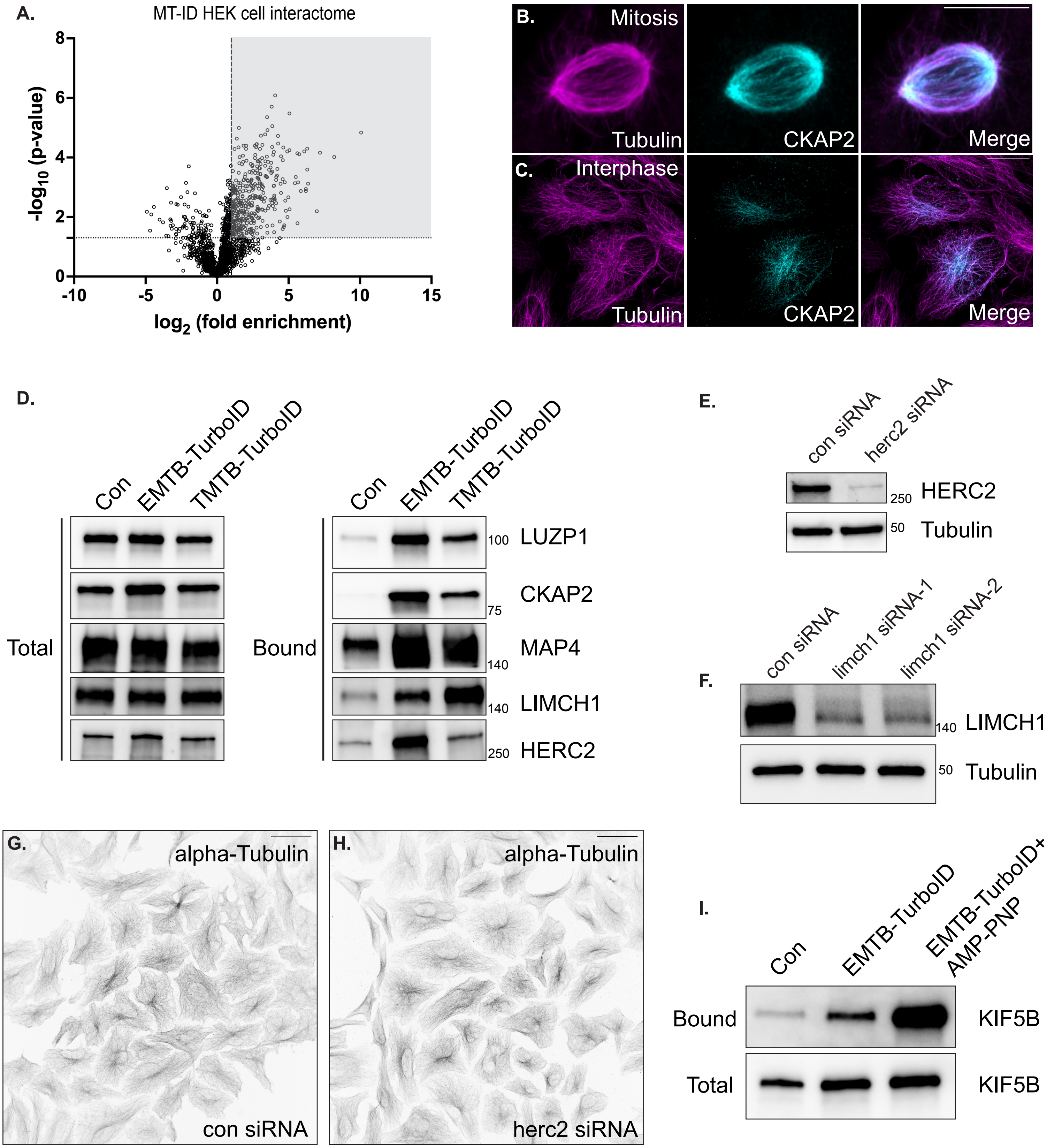


Supplemental Fig. 2

### Supplemental Figure3

A.

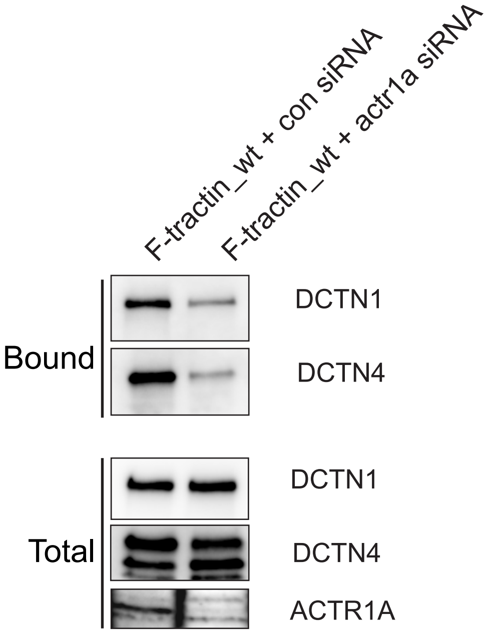


B.

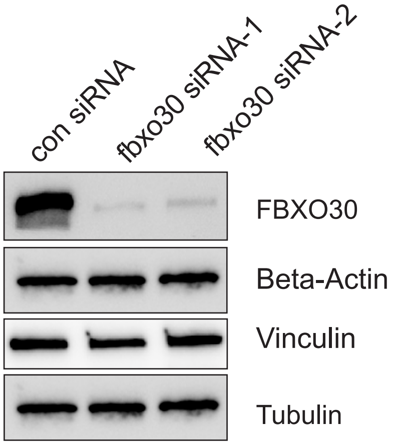


Supplemental Fig. 3
